# Comparison of fitness effects in the earthworm *Eisenia fetida* after exposure to single or multiple anthropogenic pollutants

**DOI:** 10.1101/2022.02.17.480840

**Authors:** Anja Holzinger, Magdalena M. Mair, Darleen Lücker, Dimitri Seidenath, Thorsten Opel, Nico Langhof, Oliver Otti, Heike Feldhaar

**Author notes:** **Corresponding author:** Magdalena M. Mair. These authors contributed equally (Anja Holzinger, Magdalena M. Mair). These authors contributed equally (Oliver Otti, Heike Feldhaar).

## Abstract

Terrestrial ecosystems are exposed to many anthropogenic pollutants. Non-target effects of pesticides and fertilizers have put agricultural intensification in the focus as a driver for biodiversity loss. However, other pollutants, such as heavy metals, particulate matter, or microplastic also enter the environment, e.g. via traffic and industrial activities in urban areas. As soil acts as a potential sink for such pollutants, soil invertebrates like earthworms may be particularly affected by them. Under natural conditions soil invertebrates will likely be confronted with combinations of pollutants simultaneously, which may result in stronger negative effects if pollutants act synergistically.

Within this work we study how multiple pollutants affect the soil-dwelling, substrate feeding earthworm *Eisenia fetida*. We compared the effects of the single stressors, polystyrene microplastic fragments, polystyrene fibers, brake dust and soot, with the combined effect of these pollutants when applied as a mixture. Endpoints measured were survival, growth, reproductive fitness, and changes in three oxidative stress markers. We found that among single pollutant treatments, brake dust imposed the strongest negative effects on earthworms in all measured endpoints including increased mortality rates. Sub-lethal effects were found for all pollutants. Exposing earthworms to all four pollutants simultaneously led to effects on mortality and oxidative stress markers that were smaller than expected by the respective null models. These antagonistic effects are likely a result of the adsorption of toxic substances found in brake dust to the other pollutants. With this study we show that effects of combinations of pollutants cannot necessarily be predicted from their individual effects and that combined effects will likely depend on identity and concentration of the pollutants.

## Introduction

Terrestrial ecosystems, including the soil-dwelling fauna, are exposed to many anthropogenic stressors such as an increasing number of pollutants in addition to rising temperatures or droughts due to climate change (Rodriguez-Eugenio et al. 2018). Investigating how human activities affect soil organisms is crucial because these organisms contribute to many essential ecosystem functions, such as decomposition and nutrient cycling (Blouin et al. 2013; Geisen et al. 2019; Lavelle et al. 2006). In order to effectively protect organisms in a contaminated environment, management requires knowledge of the effects of pollutants on organisms. Pollutants can originate from traffic, industrial production and agriculture (Cachada et al. 2018; De Silva et al. 2021; Gunstone et al. 2021; Rodríguez-Eugenio et al. 2018). They enter the environment via deliberate application or leakage and poor waste management (Briggs 2003). Because agricultural intensification is one of the most apparent reasons for the observed changes in biodiversity, research has focused on the side-effects of pesticides and fertilizer application on non-target organisms (Gunstone et al. 2021; Sanchez-Bayo and Wyckhuys 2019). However, especially in or close to urban areas, industrial pollution comprising heavy metals, airborne particulate matter, or microplastic deriving from degraded plastic waste released into the environment may also adversely affect invertebrate populations (Büks et al. 2020; De Silva et al. 2021; Feldhaar and Otti 2020; Ji et al. 2021; Santorufo et al. 2012b). A potential sink for such pollutants is soil, as contaminants can accumulate therein over long periods (Cachada et al. 2018; Corradini et al. 2019). As a result, soils may contain mixtures of pollutants originating from various anthropogenic activities over the years. Such mixtures could be very problematic for soil invertebrates, like earthworms, nematodes or springtails. Soil pollutants might affect soil organisms directly via toxic effects of heavy metals or pesticides (Gunstone et al. 2021; Hackenberger et al. 2018b; Pelosi et al. 2014), or indirectly by changes in the physico-chemical properties of soils such as pH or water-holding capacity or by changes in the soil microbial community (Brtnicky et al. 2021; Hackenberger et al. 2018a; Lemmel et al. 2019; Rillig et al. 2019). Rillig et al. (2019) recently showed that when more stressors (including drought and different pollutants) are combined, alterations in soil properties and microbial communities increase. However, the magnitude of these changes could not be predicted from the effects of single factors.

Studies with single pollutants can provide some information on their effects on organisms, but under natural conditions, organisms are usually exposed to a mixture of several pollutants at once. Because multiple stressors may interact in different ways, i.e. additively, synergistically or antagonistically (Piggott et al. 2015; Schäfer and Piggott 2018), estimating these effects in realistic setups is challenging. Therefore, we need studies using multiple stressors for realistic hazard estimation. Unfortunately, no general protocol exists on how to investigate multiple stressor effects experimentally. In addition, most multiple stressor studies are limited to two stressors and are not concerned with concentration dependence as a potentially confounding factor (but see Hackenberger et al. 2018b).

In recent years, studies on the effects of pollutants on soil invertebrates have focused largely on pesticides (Gunstone et al. 2021; Pelosi et al. 2014) and heavy metals (Li et al. 2020; Liu et al. 2020; Nikolaeva et al. 2019). Currently, other pollutants such as microplastic or airborne particulate matter deriving from industrial and urban sources are increasingly drawing attention from scientists due to their ubiquity and amounts released into the environment (Horton et al. 2017; Machado et al. 2018; Rillig and Lehmann 2020; Wu et al. 2019; Zhang et al. 2021a; Zhang et al. 2021b).

Soil pollution with microplastic, i.e. plastic fragments or fibers smaller than 5 mm in diameter or length respectively, and the potential entailing effects on organisms are increasingly regarded as problematic due to the amounts of microplastic entering the environment (Büks et al. 2020; Ji et al. 2021; Wang et al. 2020). Plastic enters the environment by littering, application of sewage sludge or compost, or plastic mulching as a widespread practice in agriculture (Wang et al. 2021; Weithmann et al. 2018; Wu et al. 2019). Larger plastic items are weathered and eventually fragment into microplastic due to UV exposure, mechanical stress or processing by soil organisms (Ya et al. 2021; Zhang et al. 2021a). However, microplastic can also derive directly from plastic products due to fragmentation and mechanical stress such as large amounts of tire abrasion close to streets with heavy traffic (Sheng et al. 2021; Sommer et al. 2018), or microplastic fibers shed from synthetic textiles during washing (Carney Almroth et al. 2018). Fibers and fragments are the most abundant microplastic types found in soils (Büks and Kaupenjohann 2020; Corradini et al. 2019). Soil organisms exposed to microplastics may suffer lethal and sublethal effects (Büks et al. 2020; Ji et al. 2021). Sublethal effects comprise for example increased levels of oxidative stress and damaged gut tissue, lower growth rate, and lower reproductive output (Boots et al. 2019; Chen et al. 2020; Huerta Lwanga et al. 2016; Rodriguez-Seijo et al. 2017; Rodriguez-Seijo et al. 2018; Selonen et al. 2020; Wang et al. 2019). Recent reviews suggest that the effects of microplastic on organisms depend on polymer type, size, shape, and concentration (Büks et al. 2020; Ji et al. 2021).

Airborne fine particulate matter sedimenting on the soil surface is a mostly neglected pollutant to which soil invertebrates are exposed. It derives from traffic in urban areas, agricultural dust, coal mining, and other industries (Bell et al. 2011; Kelly and Fussell 2012). Its chemical composition and size distribution strongly depends on the source of the particles. While soot dust from coal mines is mostly comprised of carbon particles, car exhaust usually comprises aggregates of a carbon core where organic residues and heavy metals are deposited (De Silva et al. 2021). Brake dust particles may contain metallic components as well as phenolic compounds from brake pads (De Silva et al. 2021; Kelly and Fussell 2012). Brake dust and soot emitted by cars can accumulate in soils near streets such as green embankments. Therefore, roadside soils can contain high levels of heavy metals such as zinc, cadmium, copper, and other harmful substances like polycyclic aromatic hydrocarbons (Nikolaeva et al. 2019; Werkenthin et al. 2014), with detrimental effects on invertebrate soil biota (De Silva et al. 2021; Nikolaeva et al. 2019).

In this study we investigate how multiple pollutants affect the soil-dwelling, substrate feeding earthworm *Eisenia fetida*. Earthworms are essential organisms in decomposition and nutrient cycling in soil (Blouin et al. 2013). Besides, they are unspecific substrate feeders that are likely to ingest pollutants contained in the soil and will come into contact with pollutants with their skin. In the present study, we compared the effects of single stressors, i.e. microplastic fragments, microplastic fibers, brake dust and soot, with the combined effect of these pollutants when applied as a mixture. We hypothesize that the mixture would lead to more severe effects than each stressor separately because of a generally increased stress response. In addition, we compare the effect of the mixture with the expected effect resulting from an additive effect model to investigate whether the pollutants act as synergists, antagonists or in an independent additive way.

## Material & Methods

### Earthworm culture

*Eisenia fetida* earthworms used in this study were reared in wormeries (Wurmfarm 42×42×60 cm, vidaXL, Venlo, Netherlands) at 15 °C and 77 % humidity in a climate chamber with a 12 h:12 h light and dark photoperiod at the Department of Animal Ecology I at the University of Bayreuth. The worms were cultivated in a mixture of 3 kg potting soil (Blumenerde, Floragrad, Bayreuth, Germany), 650 g coconut fibers (Humusziegel.de, Wörmlitz, Germany) and 300 g sphagnum peat (Floratorf, Floragrad, Bayreuth, Germany). The humidity of the soil was periodically adjusted by adding around 400 ml of deionized water weekly. Once a week, the worms were given three tablespoons of oatmeal and a tablespoon of special worm food (Superwurm e.K., Düren, Germany). Both foods were evenly distributed on top of the soil in the wormery. At the start of the experiment, all individuals had no clitellum and were of similar weight (250 to 400 mg).

### Anthropogenic pollutants

#### Polystyrene (PS) fragments and fibers

We chose polystyrene (PS) as polymer type to study the effects of microplastic. PS is among the most commonly used polymer types used for production of plastic products (PlasticsEurope 2020). Granules and fibers were ordered from Styrolution (PS 158N/L, Frankfurt am Main, Germany) and further processed to fragments and fibers by the Department of Macromolecular Chemistry I (MCI) at the University of Bayreuth. Polystyrene granules were ground to fragments using a cryo ball mill (Retsch, CryoMill, Germany) followed by sieving to obtain fragments ranging from 125-200 μm in diameter. Fibers were milled in a laboratory blender (Waring LB20ES, Beechwood, United Kingdom) for 30 minutes to achieve a length of approximately 1-4 mm and a diameter of 40 µm.

#### Brake dust particles

Brake dust particles were provided by the Department of Ceramic Materials Engineering of the University of Bayreuth. Tribologically tested LowMet brake pads (provided by TMD company), a pad type which is applied for the most passenger cars in europe, were ground, after several braking cycles on a ceramic brake disc, i. e. after a dissipation of a total friction energy of about 15 MJ and after reaching temperatures up to 400 °C. The full braking cycles were conducted on ceramic brake discs with a diameter of about 400 mm and initial sliding speeds between 5 m/s and 20 m/s. The brake pads were ground for three minutes in a vibrating cup mill with a tungsten carbide grinding set (Pulverisette 9, Fritsch GmbH, Idar-Oberstein, Germany) to reach a fine-grained powder. The particle size distribution of the ground brake pads was measured using a laser diffraction particle size analyzer (PSA 1190 LD, Anton Paar GmbH, Ostfildern-Scharnhausen, Germany). The average particle size found was 10.19 ± 4.37 µm mean ± SD (D10 = 0.68 µm (10% of all particles being smaller in diameter than this size), D50 = 5.76 µm (median particle size), D90 = 25.87 µm (90% of particles being smaller in diameter than this size). LowMet brake pads consist of steel wool (15 % (w/w)), petrol coke (12 % (w/w)), sulphides (10 % (w/w)) as well as aluminum oxide and binder (both 5 % (w/w)) as largest fractions (Wiaterek 2012). Binders and fillers comprise metallic and phenolic compounds (Chan and Stachowiak 2004).

#### Soot particles

We used the carbon black PRINTEX 30 Furnace Black (Degussa AG, Frankfurt, Germany) for the soot treatments with an average primary particle size of 27 nm. Carbon black and soot are often used interchangeably even though they are distinct from each other. Carbon blacks are commercially produced elemental carbon particles with different but defined physico-chemical properties (Watson and Valberg 2001). In contrast, soot is a by-product of relatively uncontrolled, incomplete combustions, which results in a material of varying and often unknown composition, but with particles of similar size between soot and carbon blacks (Clague et al. 1999). In this study we used carbon black as a surrogate for diesel engine soot to simulate contaminated soil. Soot is the most similar, naturally occurring pendant, we henceforth refer to the carbon black as soot.

#### Exposure of *E*.*fetida* to pollutants

We used individual juvenile earthworms (i.e. without clitellum) with a bodyweight between 250 mg and 400 mg for the experiment. The juveniles were of unknown age. We placed the selected earthworms in plain test soil for one week under the same conditions as described above for the laboratory culture to adjust to the new environment. We then performed the exposure assay according to the OECD guideline 222 for the earthworm reproduction test (OECD 2016). The test substrate was a mixture of 70 % quartz sand, 20 % kaolin clay (Gebr. Dorfner, Hirschau, Germany) and 10 % sphagnum peat. We air-dried and mixed the components thoroughly. Soil pH was adjusted to pH 5.97 by adding calcium carbonate. The moisture content of the soil used in the experiment corresponded to 60% of the maximum water holding capacity, which was determined using the method according to the standard ISO DIS 11268-2 (https://www.iso.org/standard/53528.html).

As test vials, we used 800ml glass jars (WECK, Wehr, Germanys). Plain polystyrene fragments and fibers, brake dust and soot were added individually to premixed dry OECD soil at 0.5 % (v/v) and 2 % (v/v) to a final weight of 100g of soil mixed with pollutants. In addition, we prepared three concentrations of multiple pollutant treatments, i.e. 0.5 % (v/v), 2 % (v/v) and 8 % (v/v). Therefore, we mixed all four pollutants at equal proportion (v/v). By using the three different concentrations of multiple pollutants we wanted to test whether the mixture of pollutants affected the earthworm differently in comparison to the same volume of a single pollutant or whether the higher volume of pollutants affected the worms more when pollutants were combined. As a control treatment, we used plain test substrate without any pollutants. For each replicate, we then randomly selected six earthworms, weighed to the nearest 0.01 g (KERN PFB, Kern & Sohn GmbH, Balingen, Germany). Then we transferred them to a glass jar on top of the treated soil substrate. For each treatment, we prepared eight replicate glass jars. The order in which treatments were prepared was randomized.

The worms were exposed to treatments for eight weeks. Once a week the worms received 0.5 g of special worm food (Superwurm e.K., Düren, Germany). A thin layer of food was spread evenly on the ground and moistened with three sprays of deionized water from a spray bottle. Using deionized water prevents the food from going moldy but maintains the moisture. After four weeks of exposure, we removed the worms from each jar separately, rinsed them carefully with distilled water, counted the number of dead and living worms, weighed the live worms, and then put them back into their jars. Worms were considered dead when they were either missing or did not move any longer when handled. After eight weeks we weighed and counted them again in the same manner and then randomly selected three worms from each jar to measure the oxidative stress response and lipid peroxidation. The randomly selected individuals were transferred to a petri dish with a moist filter paper and kept in the climate chamber for 24h to cleanse their gut contents by defecation. After this, we snap froze these individuals in liquid nitrogen until performing the stress response assays (see Effect of exposure of pollutants on the oxidative stress response and lipid peroxidation).

### Reproductive output

To measure the reproductive output, we counted the number of cocoons and juveniles per glass jar. Hence, we placed the jars in the climate chamber for an additional two weeks to allow the juveniles to hatch after the removal of all adult individuals (see above). Thus, after a total of ten weeks after the start of the experiment, we placed the glass jars in a water bath at 30 °C and slowly increased the water temperature to 50 °C. After about 20 minutes, the juvenile worms started to appear on the soil surface and were collected. Then we counted and stored the juveniles in a freezer at -80 °C until further use. To determine the number of cocoons, we emptied each glass jar separately, spread the soil substrate in a flat tray, and visually screened the substrate and counted all cocoons.

### Oxidative stress response and lipid peroxidation

Oxidative stress induced by reactive oxygen species (ROS) can arise either due to an oxidative challenge directly created by a pollutant or indirectly as a by-product of increased aerobic respiration (Trestrail et al. 2020). The antioxidant response of earthworms was evaluated by measuring the activity of the two oxidative stress-related biomarkers, glutathione S-transferase (GST) and catalase (CAT). GST is an antioxidant enzyme that directly neutralizes ROS and detoxifies pollutants (Aly and Schröder 2008; LaCourse et al. 2009; Trestrail et al. 2020). Catalase (CAT) is an important enzyme involved in removing hydrogen peroxide (H_2_O_2_). We assessed the GST activity according to Habig et al. (1974) and the CAT activity using the method as described in Hadwan (2018). In addition, we measured lipid peroxidation by quantifying the biomarker malondialdehyde (MDA) content as described in Buege and Aust (1978).

The whole frozen worms collected after eight weeks of exposure were homogenized by hand with a plastic mortar in 700µl phosphate buffer on ice (150 mM, pH = 7.0 mixed with 0.1 % Triton X-100). The homogenates were centrifuged at 10,000 g and 4 °C for 10 minutes. The supernatant was transferred to a fresh 1.5 ml Eppendorf tube and stored at -20 °C until analyzed. All three stress responses were measured in a spectrophotometer equipped with a microplate reader (Synergy HT5, BioTek, Bad Friedrichshall, Germany). All biomarkers were expressed relative to the total protein content for each sample, which was determined with a Bradford assay (Bradford 1976) using the Bradford reagent (Carl Roth, Karlsruhe, Germany). Results were expressed as enzymatic activity per mg protein for CAT and MDA and as enzymatic activity in nM per minute per mg protein content for GST.

### Statistical analysis

All analyses were done in R version 4.1.0 (R Core Team 2021). Growth models were fitted using the R package *metafor* version 3.0-2 (Viechtbauer 2010). All other models were fitted using *glmmTMB* version 1.1.2 (Brooks et al. 2017). Residuals were checked using *DHARMa* version 0.4.3 (Hartig 2021).

For all measured endpoints, we tested whether the exposure to the different pollutants changed the outcome compared to the control treatment. In all models, treatment was used as a fixed factor with 12 levels: control, low and high concentration of brake dust, low and high concentration of soot, low and high concentration of microplastic fragments, low and high concentration of microplastic fibers, and low, medium and high concentration of a mix of all four pollutants.

#### Exposure effects on survival

Survival was analyzed by fitting a generalized linear mixed model (glmm) with binomial error structure and logit link. As six worms were exposed together in a glass jar, we added a random intercept for the unique glass IDs. The survival status after four and eight weeks of exposure was analyzed separately. A single surviving individual was added to treatments where all worms had died to avoid zero variance and complete separation, which results in unreasonably large standard errors and p-values.

#### Exposure effect on growth

Bodyweight (measured in g) was analyzed by fitting a linear model with known sampling variances to the mean bodyweight of earthworms per unique glass jar. The analysis was based on means and sampling variances per glass jar, because the same earthworms were measured repeatedly (before exposure, after four and after eight weeks of exposure), but their identities could not be tracked. We tested whether the initial weight of earthworms in all treatments containing pollutants was the same as in the control treatment. An interaction term between treatment and exposure time (week) was added to test whether the change in bodyweight over eight weeks (the slope of weight over time, i.e. growth) differed between the control and the exposure treatments.

#### Exposure effect on the reproductive output

Reproductive output was analyzed by fitting two separate generalized linear models (glm) with negative binomial error structure and log-link to the number of cocoons and the number of juveniles found in glass jars after ten weeks of exposure, respectively. A single cocoon (or juvenile) was added to treatments with all-zero counts to avoid issues with complete separation. Because offspring counts were done per glass jar, no random effect was added.

#### Exposure effect on the oxidative stress response and lipid peroxidation

The enzymatic activity of catalase (CAT) in units per mg protein was analyzed by fitting a generalized linear mixed model with gamma-distributed error term and log-link. The enzymatic activity of glutathione S-transferase (GST) in nmol per minute per mg protein content and malondialdehyde (MDA) in units per mg protein as a marker for lipid peroxidation were analyzed with linear mixed models (lmm). The unique glass ID was again fitted as a random effect in all three models.

#### Expected additive effects

Effects expected for the medium and high mixed pollutant treatments were calculated based on the predicted effects in the single pollutant treatments (0.5 and 2 v/v%, respectively). We used different null models for different endpoints based on recommendations in Schäfer & Piggott (2018): expected effects for survival were calculated using a multiplicative null model; because effects of brake dust on reproduction were close to the effect limit (zero offspring), we used the dominance null model for the calculation of the expected number of cocoons and juveniles in the mix treatments; for growth, oxidative stress response and lipid peroxidation, simple addition was used as the null model for the expected multiple pollutant effects.

#### Open data and code

All raw data and R code pertaining to the manuscript are available at https://github.com/magdalenammair/Holzinger-et-al-2022 and https://doi.org/xx.xxxx/zenodo.xxxx.

## Results

### Exposure effect on survival

After four weeks of exposure, survival rates of worms exposed to brake dust were lower than for worms in the control treatment at both low (glmm, z = 3.68, p = 0.0002; Figure 1A) and high concentrations (glmm, z = 5.74, p < 0.0001). Irrespective of particle concentrations neither exposure to soot, nor microplastic fragments or fibers affected earthworm survival (glmm, all p-values > 0.1, see Table S1). Earthworms exposed to a mix of all four pollutants were negatively affected only in the high concentration treatment (glmm, z = 4.96, p < 0.0001), but not at low (glmm, z = 1.18, p = 0.24) or medium concentrations (glmm, z = -0.00, p = 0.99). The survival rate in the medium and high concentration mix treatment was higher than expected based on a multiplicative effect model (see red triangles in Figure 1A).

**Figure 1.**
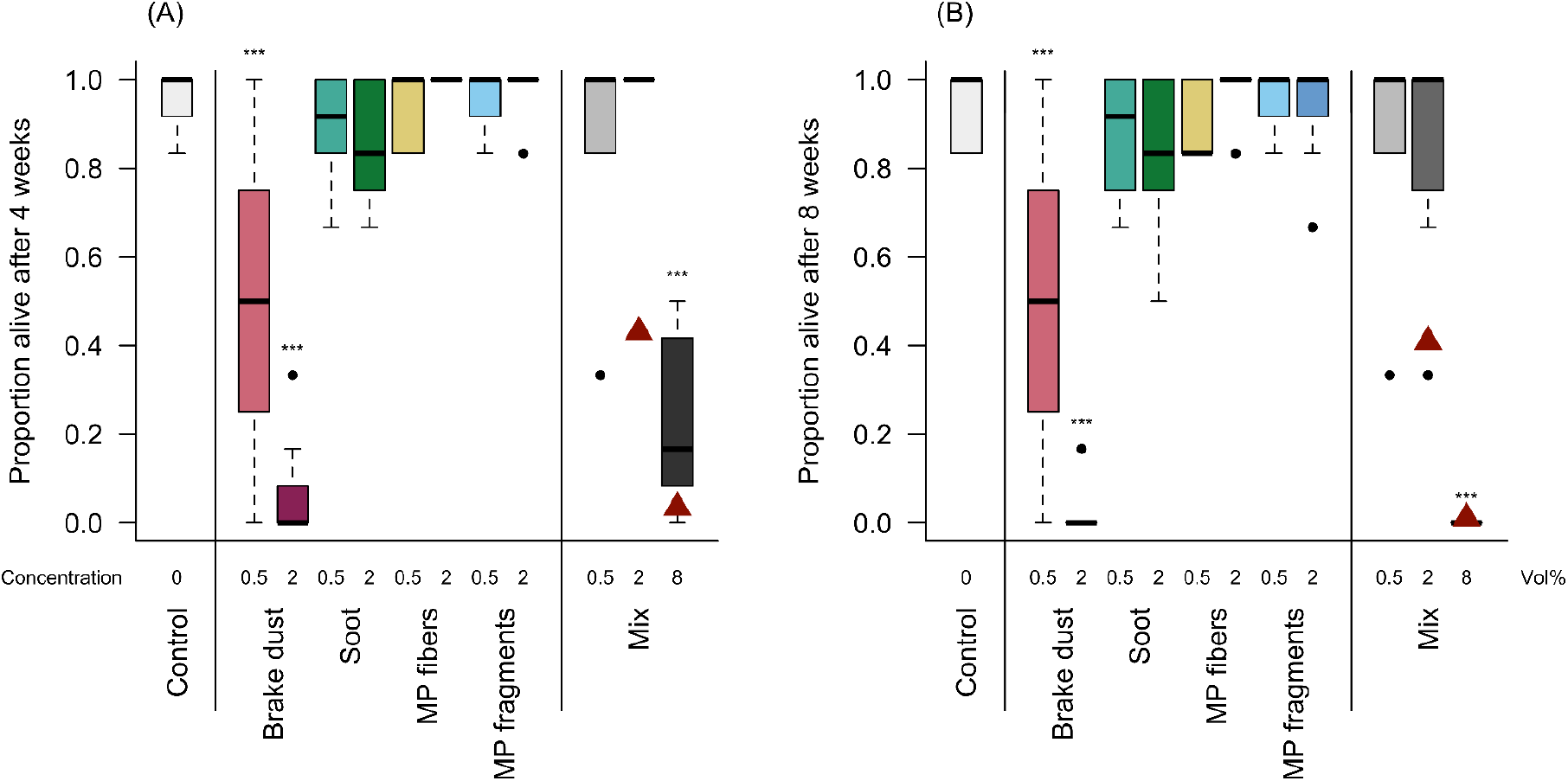
Proportion of earthworms alive in glass jars after (A) four and (B) eight weeks of exposure to different pollutants. Asterisks indicate significant differences to the control treatment (logistic regression; *** p < 0.001). Red triangles: expected effects (multiplicative model).

The pattern in survival rates was almost identical after eight weeks of exposure (Figure 1B): Reduced survival was observed in treatments with low (glmm, z = 3.42, p = 0.0006) and high (glmm, z = 5.36, p < 0.0001) concentrations of brake dust and the high concentration mix treatment (glmm, z = 5.40, p < 0.0001). No negative effects on survival were observed in treatments with soot, microplastic fibers, microplastic fragments and the low and medium mix treatments (glmm, all p-values for comparisons with the control treatment > 0.2, see Table S2). After eight weeks, the survival rate in the medium concentration mix treatment was higher than expected by the multiplicative effect model (see red triangles in Figure 1B). As expected by the multiplicative model, in the high concentration mix, none of the worms survived to week eight.

### Exposure effect on growth

Earthworm bodyweight was the same in all treatments at the beginning of the experiment (Figure 2A, before exposure; linear model, all p-values for comparisons with the control treatment > 0.1; see Table S3). Over the course of the experiment, brake dust negatively affected earthworm growth at both low (lm, z = -6.67, p < 0.0001) and high concentrations (lm, z = - 5.30, p < 0.0001): Earthworms exposed to low concentrations of brake dust did not grow over eight weeks, and worms exposed to high concentrations of brake dust even lost weight. Exposure to a mix of all four pollutants did not affect earthworm growth at low concentrations (lm, z = 0.38, p = 0.71), and led to weight loss at medium (lm, z = -10.31, p < 0.0001) and high concentrations (lm, z = -12.33, p < 0.0001) (Figure 2B, C). Earthworms in all other treatments showed similar growth to the growth in the control treatment (p-values for all interactions > 0.1; see Table S3). After eight weeks of exposure, all worms in the high concentration mix treatment were dead and no weight measurements were taken. Growth in the mix treatments did not differ from effects expected from a simple addition null model (see red triangles in Figure 2 illustrating expected weights in week 4 and 8 calculated based on expected growth from the simple addition null model).

**Figure 2.**
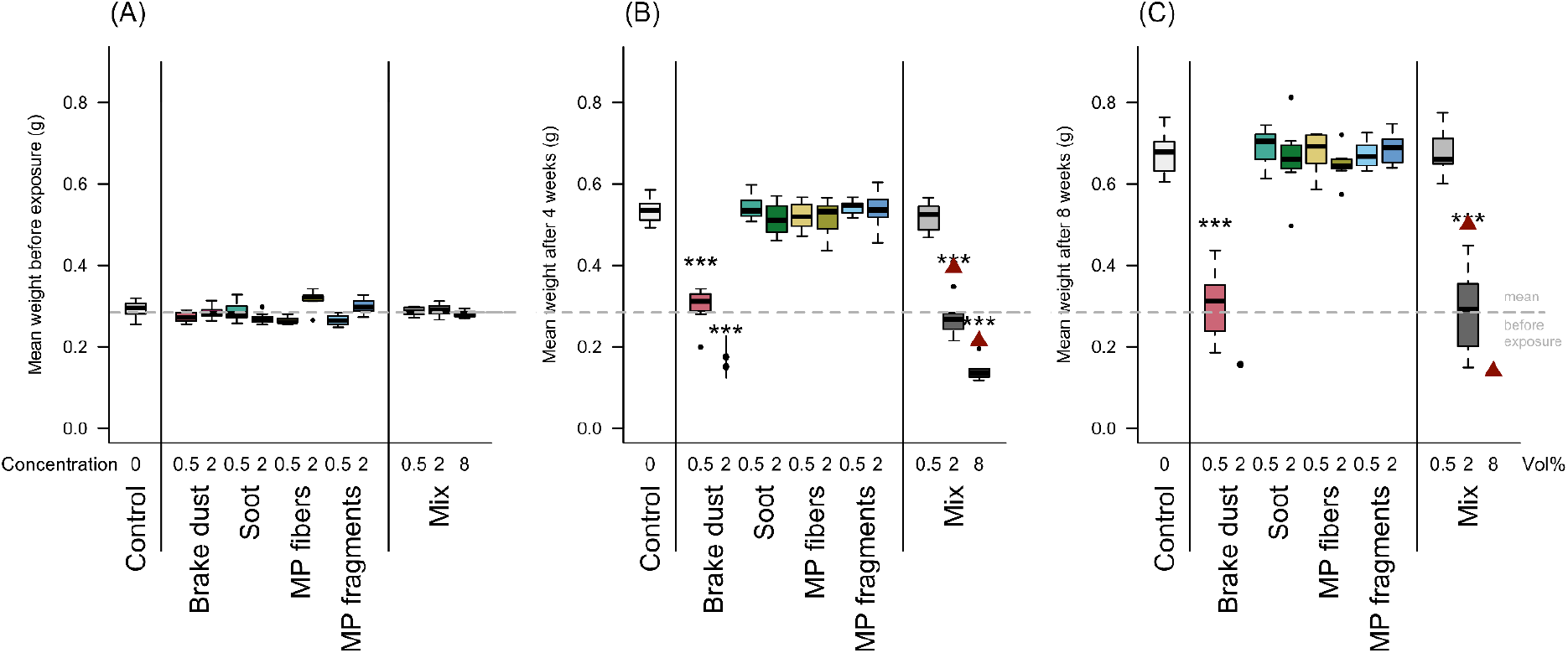
Bodyweight in g at (A) the start of the experiment, (B) after four weeks and (C) eight weeks of exposure to different pollutants. Asterisks indicate significant differences to the control treatment (logistic regression; *** p < 0.001). Grey dashed line: control treatment mean before exposure. Red triangles: expected effects (simple addition null model).

### Exposure effect on the reproductive output

The number of cocoons and juveniles found in substrates after ten weeks of exposure (adult earthworms removed after 8 weeks) was reduced in treatments with soot at low (glmm, cocoons: z = -4.99, p < 0.0001 ; juveniles: z = -5.18, p < 0.0001; Figure 3) and high concentrations (glmm, cocoons: z = -5.44, p < 0.0001 ; juveniles: z = -6.04, p < 0.0001) and with low concentrations of microplastic fibers (glmm, cocoons: z = -5.86, p < 0.0001 ; juveniles: z = -6.56, p < 0.0001) and low concentrations of microplastic fragments (glmm, cocoons: z =-5.03, p < 0.0001 ; juveniles: z = -5.18, p < 0.0001). No offspring was produced by earthworms exposed to brake dust (glmm, cocoons: low and high concentration: z = -5.84, p < 0.0001; juveniles: low and concentration: z = -5.18, p < 0.0001; for comparisons with the control treatment). No negative effects on offspring numbers were observed in treatments with high concentrations of microplastic fibers and fragments (glmm, cocoons and juveniles: all p-values for comparisons with the control treatment > 0.3; see Tables S4 and S5).

**Figure 3.**
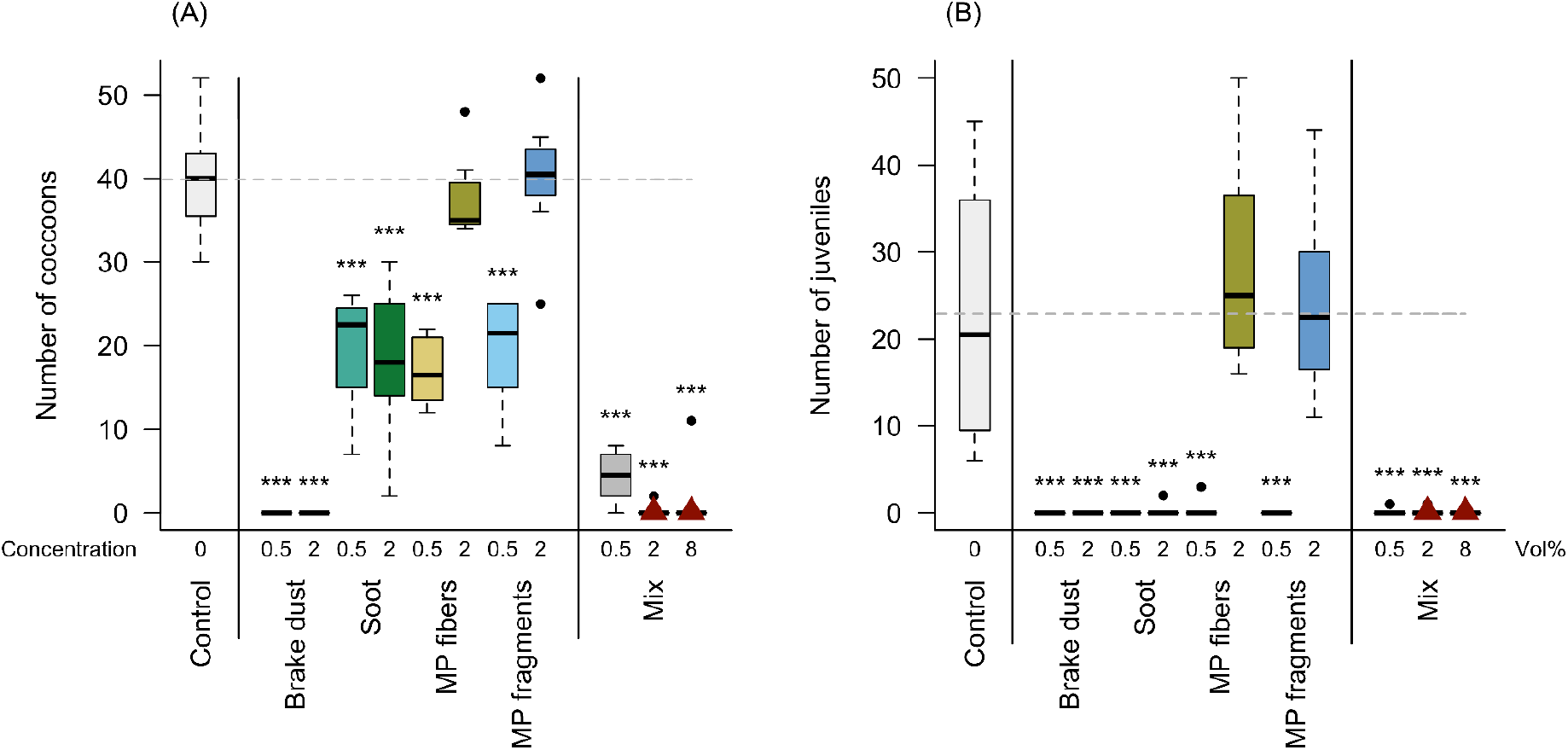
Number of (A) cocoons and (B) juveniles found in the substrate after 10 weeks of exposure to different pollutants. Adult parents have been removed from the substrate after 8 weeks. Asterisks indicate significant differences to the control treatment (glmm with negative binomial error distribution): ***: p < 0.001. Grey dashed line: control treatment mean. Red triangles: expected effects (dominance model).

### Reproductive output in the mix treatments was in general low

While worms in the low concentration mix treatment produced some cocoons (see Figure 3A), the cocoons were almost entirely absent in the medium and high mix treatments (cocoons found in only one jar each). Juveniles were only found in one jar of the low and medium mix treatment, respectively (Figure 3B). These low reproduction rates were in accordance with the expected effects from the dominance null model.

### Exposure effect on the oxidative stress response and lipid peroxidation

Because all worms had died after eight weeks of exposure in the high concentration brake dust and high concentration mix treatment, no radical oxygen species (CAT and GST) and lipid peroxidation measurements (MDA) were taken for these treatments.

CAT activity was increased in surviving worms exposed to low concentrations of brake dust (glmm, z = 3.26, p = 0.001; Figure 4A), low concentrations of soot (glmm, z = 2.01, p = 0.04), high concentrations of microplastic particles (glmm, z = 2.45, p = 0.01) and low (glmm, z = 2.98, p = 0.003) and medium concentrations (glmm, z = 2.21, p = 0.03) of the pollutant mix.

**Figure 4.**
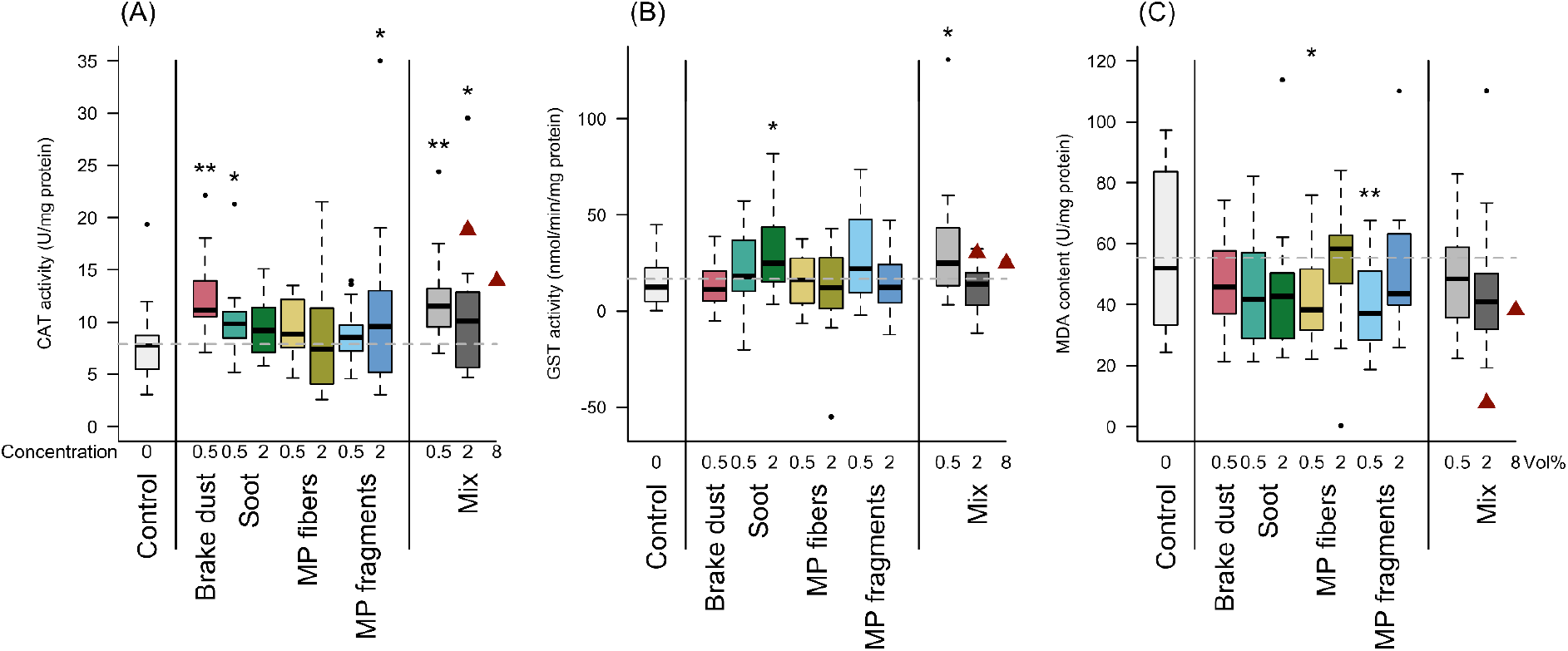
Enzymatic activity of (A) catalase (CAT) in units per mg protein, (B) glutathione S-transferase (GST) in nmol per minute per mg protein content, and (C) malondialdehyde content (MDA) in units per mg protein in earthworms that survived the exposure to different pollutants for eight weeks. No measurements are available for the high concentration of brake dust and high concentration mix treatment because all worms died in these treatments. MP stands for microplastic. Asterisks indicate significant differences to the control treatment (CAT: generalized linear mixed model with gamma distributed error term and log link, GST and LPO: linear mixed model): * p < 0.05, ** p < 0.01. Grey dashed line: control treatment mean. Red triangles: expected effects (simple addition null model).

Soot at high concentrations, microplastic fibers and microplastic particles did not affect CAT activity (glmm, all p-values for comparisons with control > 0.1; see Table S6.

GST activity was only increased in earthworms exposed to high concentrations of soot (linear mixed model, z = 2.41, p = 0.02; Figure 4B) or low concentrations of mixed pollutants (lmm, z = 2.31, p = 0.02). All other treatments did not show a significant increase in GST activity (lmm, all p-values for comparisons with control > 0.06; see Table S7).

MDA activity was decreased in earthworms exposed to low concentrations of microplastic fibers (linear mixed model, z = -2.08, p = 0.04; Figure 4C) and low concentrations of microplastic particles (lmm, z = -2.72, p = 0.007). No effects on lipid peroxidation were observed in earthworms in the other treatments (lmm, all p-values for comparisons with control > 0.09; see Table S8).

In all three measured enzymes, mean effect sizes in the medium mix treatments were smaller than expected by the simple addition null model (see red triangles in Figure 4A-C). As no worms survived to week eight in the high concentration mix treatment, a comparison between measured and expected effects was not possible (expected effects are nevertheless shown in Figure 4).

## Discussion

In this study, we exposed earthworms to the four single pollutants: polystyrene (PS) microplastic fragments, PS fibers, soot, and brake dust at two different concentrations and the combination of all four pollutants in three different concentrations to determine dose-dependent effects and interactions between these pollutants. We found that soil pollution affects different fitness parameters and stress markers in *Eisenia fetida* depending on the pollutant type and in a multiple stressor scenario on the concentration.

Brake dust was by far the most detrimental of the four pollutants: mortality of earthworms was significantly increased (Fig. 1) and growth was significantly slower (Fig. 2). In contrast, the other three single pollutants neither affected mortality nor growth. A reason why brake dust might be harmful to *E. fetida* is that it contains potentially toxic metals (such as antimone, barium, molybdenum, copper or zinc) and phenolic compounds (Chan and Stachowiak 2004; Gietl et al. 2010; Volta et al. 2020). However, the mechanism how negative effects are mediated is not entirely clear yet and requires further studies. In single pollutant acute contact toxicity tests using only brake dust (comparable to the brake dust used in our study), Volta et al. (2020) in contrast did not find negative fitness effects on the earthworm *Eisenia andrei*. The difference to our experimental setup however was that Volta et al. (2020) exposed earthworms to brake dust suspensions on filter paper for 48h instead of in-soil exposure for several weeks, and used concentrations that were approximately one order of magnitude lower (highest concentration: 1ml of a solution containing 1000 mg of brake dust/L; i.e. 0.1 % (w/w)) than the concentrations we used (0.5 % (v/v) which equals 1.04 % (w/w) in the low concentration treatment). By adding brake dust directly to the soil substrate, worms in our study not only came into contact with brake dust with their body surface, but also ingested the potentially toxic substance during feeding in a chronic toxicity test setup. Negative impacts of roadside soils likely containing brake dust on earthworm survival and growth have recently been shown (Nikolaeva et al. 2019). However, these soils were not only contaminated with brake dust but also other pollutants such as polycyclic aromatic hydrocarbons. A study using soils from sites close to roadsides around downtown Naples for similar bioassays as used here showeds sublethal effects on growth and reproduction of *E. fetida*, but no lethal effects (Santorufo et al. 2012a). Taken together, our finding of strong negative fitness effects on earthworms kept in artificial soil polluted with brake dust fits well to the negative fitness effects found in the studies using roadside soils (Santorufo et al. 2012a; Nikoleava et al. 2019), which strongly suggests that brake dust indeed affects soil organisms close to roadsides.

While mortality was only increased in treatments containing brake dust, sublethal effects such as a reduction in reproductive output (number of cocoons and juveniles) and elevated levels of stress markers were observed for each of the four single pollutants, albeit not at both concentrations (Fig. 3, Fig. 4). Again, brake dust had the strongest negative effect on reproductive output as earthworms exposed to this pollutant did not produce any offspring, neither at high nor at low concentrations. Soot significantly reduced the number of cocoons as well as the number of juveniles produced at both concentrations. In contrast, in treatments with PS microplastic fragments and fibers the number of cocoons as well as juvenile worms was only significantly reduced at low concentrations but remained similar to the control at high concentration. The high level of reproductive output of the high concentration microplastic groups may be facilitated by the slightly above average body weight at the beginning of the experiment (Fig. 2A), likely resulting in an earlier onset of reproduction in these earthworms. The occurrence of cocoons and absence of juveniles in soot treatments and low concentration PS microplastic treatments (fibers and fragments) suggests that reproduction may have been delayed rather than entirely inhibited in these treatments (Fig. 3). Nevertheless, we can assume that polluted soils can produce important physiological costs for several life-history traits. Significant oxidative stress responses were also observed in all single pollutant treatments, albeit in different markers depending on the pollutant (Fig. 4). CAT activity increased in treatments with brake dust alone (low and high concentration), at high concentrations of PS microplastic fragments, and in mixed treatments. This suggests that in these treatments the production and removal of H_2_O_2_ may be increased. In contrast, Glutathione S-transferase (GST) activity was slightly but significantly increased only in earthworms exposed to high concentrations of soot as a single pollutant and the mixed treatment at the lowest concentration. In contrast to our expectation, malondialdehyde (MDA) concentration as a biomarker for oxidation levels of lipids was significantly lower in the low concentration PS microplastic fragment and fiber treatments compared. An increase in CAT activity and decrease in GST activity was also found in earthworms confronted with PS fragments by Wang et al. (2019). Earthworms confronted with tire wear particles containing heavy metals and microplastic likewise showed an increase in CAT and decrease in GST activity (albeit at very high concentrations from 5% to 20% w/w in soil) (Sheng et al. 2021). Other studies assessing oxidative stress responses of *E. fetida* towards pollutants showed inconclusive results, i.e. with an increase or decrease in CAT and GST activity in earthworms exposed to microplastic only at certain concentrations within a dataset but not others (overview in Trestrail et al. 2020; Rodriguez-Seijo et al. 2018; Li et al. 2021a). In addition, studies measuring oxidative stress responses at different time points have shown that CAT and GST activity changes over time (Li et al. 2021a; Sheng et al. 2021) Many studies found increased MDA concentrations when earthworms were exposed to different pollutants such as pesticides, heavy metals, microplastic, or tire wear particles (Hackenberger et al. 2018b; Huang et al. 2021; Sheng et al. 2021). In another study, MDA concentration was only elevated at unrealistically high concentrations (20% w/w) of polyethylene microplastics (Wang et al. 2019). Due to the complexity of the oxidative stress response and the number of different pathways involved, the interpretation of results presented here is not straightforward. Nonetheless, that all of the tested pollutants changed certain aspects of the earthworms’ oxidative stress shows that all of the pollutants pose a stress to earthworms.

The carbon black soot used as surrogate for exhaust particles should contain fewer contaminations with potentially toxic substances in comparison to soot particles collected from vehicle exhaust (Watson and Valberg 2001). Sublethal effects may therefore not be due to direct toxic effects by the carbon black used here, but indirect effects mediated via soil properties, nutrient availability, or impacts of soil or gut microbiome of the earthworms. Studies on effects of biochar (that resembles charcoal and has a high carbon content) on soils have shown that microbial biomass and / or activity is often reduced (Brtnicky et al. 2021). Likewise, negative effects on earthworms were often detected in soils treated with biochar (Brtnicky et al. 2021). Microplastic particles and fibers have been shown to result in negative fitness effects such as increased mortality, an increase in the production of stress markers, reduced reproductive output, or reduced activity in several studies (Chen et al. 2020; Cheng et al. 2021; Huerta-Lwanga et al. 2021; Jiang et al. 2020; Lahive et al. 2021; Prendergast-Miller et al. 2019; Rodriguez-Seijo et al. 2018; Rodriguez-Seijo et al. 2017).

The combination of all four pollutants in the mixture treatments showed an antagonistic effect on mortality. Fewer earthworms died than expected at medium pollutant concentration (2 % (v/v) in total with the soil containing 0.5 % (v/v) of each single pollutant) and also at the highest concentration (8 % (v/v)) after four weeks (Fig. 1). Here, mortality was significantly lower than expected and lower than the mortality for the low concentration of brake dust alone. Likewise, effects on CAT, GST and MDA activity in mixture treatments were smaller than expected by the respective null models (Fig. 4). This antagonistic effect may result from the adsorption of toxic substances present in brake dust to one or several of the other tested pollutants, and thus reducing their bioavailability. Soot may act similar to active charcoal or biochar. Both of these compounds have been shown to adsorb organic substances such a phenolics (Brtnicky et al. 2021; Zango et al. 2020), but also heavy metals (Duwiejuah et al. 2020; Koltowski and Oleszczuk 2016; Schweiker et al. 2014) that are contained in brake dust. Likewise, microplastic has also been shown to adsorb heavy metals and organic pollutants (Tourinho et al. 2019; Verla et al. 2019) due to its high surface area-to-volume ratio and potentially charged surfaces. Microplastic may subsequently act as a vector for these contaminants to terrestrial fauna and may increase (Huang et al. 2021; Li et al. 2021b; Verla et al. 2019) or reduce bioavailability (Verla et al. 2019; Yu et al. 2020).

Not all soil organisms are exposed to pollutants in the same way. Some of them only use it as shelter and breeding ground and others take up food particles from the soil surface or ingest the soil as substrate feeders. This might be a reason why we observed larger lethal and sublethal effects in *E. fetida* than we reported earlier for ants (Seidenath et al. 2021). In the earlier study we kept ant queens in soils with the same pollutant treatments used in the present experiment. We found neither negative fitness effects on the queens themselves nor on their brood (Seidenath et al. 2021). In contrast to ant queens and their brood, which only come into contact with the pollutants with their body surface, earthworms also ingest the pollutants when feeding on the substrate. Ingesting pollutants may be more detrimental as the intestinal tract is likely to be more vulnerable than the sclerotized insect cuticle. In addition, unlike ants, earthworms take up some substances from the soil through their skin, such as the potentially toxic metals copper (Cu) and lead (Pb) (Vijver et al. 2003).

While we wanted to test interactive effects when earthworms are exposed to single and a mixture of pollutants, the question arises whether the level of pollutants chosen here would be relevant under natural conditions. Hamilton and Hartnett (2013) showed that soils in urban areas of Phoenix, Arizona contained between 0.02 to 0.54 % (w/w) carbon black from anthropogenic sources. Our low and high concentration of soot falls within this range with 0.5 % (v/v) being equal to 0.08 % (w/w) and 2 % (v/v) to 0.31 % (w/w) respectively (see Table S7). Highly polluted soils in industrial areas contained up to 6.7 % (w/w) of microplastic (Büks and Kaupenjohann 2020; Chae and An 2018; Fuller and Gautam 2016), which is even higher than the concentrations we have used in the present study with the highest concentration of 2 % (v/v) being equal to 1.05 % (w/w) (see Table S7).

## Conclusions

In conclusion, we showed that exposure to single pollutants differs in the negative impact on earthworm fitness, with brake dust being by far the most harmful single pollutant in comparison to PS fragments or fibers or soot. Higher concentrations (2 % vs. 0.5 % (v/v)) of single pollutants led to stronger negative effects in some treatments but not others. The exposure to a combination of pollutants was less harmful than expected. Currently, we still know too little about the interplay of different pollutants on earthworms and other soil organisms. Interactive effects may attenuate negative effects, as was observed in the mixed pollutants treatments in the present study. Under natural conditions, animals will likely always face a combination of pollutants and potentially additional stressors such as drought which may exacerbate effects. In addition, long-term interactions among pollutants and the environment may change the pollutants’ physico-chemical properties, which may again influence their effects on soil biota. A range of pollutants may then act as vectors for toxic substances, as was shown for microplastic and soot and biochar, or may affect soil biota indirectly through changes in soil properties or interactions with other organisms such as microbiota.

## Acknowledgements

We thank Sara Pölloth, Melanie Rothmaier, Max Döring and Lisa Albert for helping with the experimental work. Furthermore, we thank Andreas Mittereder for providing us with carbon black.

## Author contributions

AH, OO, DS, DL, HF conceived the idea and planned the experiment. AH, DL, NL and TO produced the particles. DL and AH conducted the experimental work. AH and MM performed the statistical analyses. AH, OO, DL, HF and MM wrote the manuscript. All authors read and approved of the final manuscript.

