## Supplemental Tables for "Comparison of fitness effects in the earthworm *Eisenia fetida* after exposure to single or multiple anthropogenic pollutants"

---

Table S1. Regression table for the generalized linear mixed model on earthworm survival after 4 weeks of exposure to pollutants. The intercept represents the control treatment.

| | Estimate | SE | z | p |
|---|---|---|---|---|
| Intercept(Control) | -3.58 | 0.873 | -4.10 | <0.001 |
| Brake dust 0.5 | 3.58 | 1.004 | 3.56 | <0.001 |
| Brake dust 2 | 6.75 | 1.218 | 5.54 | <0.001 |
| Soot 0.5 | 1.06 | 1.058 | 1.00 | 0.315 |
| Soot 2 | 1.51 | 1.030 | 1.46 | 0.144 |
| MP fibers 0.5 | 0.46 | 1.119 | 0.41 | 0.678 |
| MP fibers 2 | -0.70 | 1.376 | -0.51 | 0.612 |
| MP fragments 0.5 | 0.00 | 1.192 | 0.00 | >0.999 |
| MP fragments 2 | -0.74 | 1.386 | -0.53 | 0.595 |
| Mix 0.5 | 1.17 | 1.051 | 1.11 | 0.267 |
| Mix 2 | -0.70 | 1.376 | -0.51 | 0.612 |
| Mix 8 | 5.02 | 1.049 | 4.79 | <0.001 |

*Notes.* Model formula: Survival ~ treatment + (1 | GlassID); Family: binomial(logit);SE: Standard error.

Table S2. Regression table for the generalized linear mixed model on earthworm survival after 8 weeks of exposure to pollutants. The intercept represents the control treatment.

| | Estimate | SE | z | p |
|---|---|---|---|---|
| Intercept(Control) | -3.13 | 0.764 | -4.09 | <0.001 |
| Brake dust 0.5 | 3.13 | 0.916 | 3.42 | <0.001 |
| Brake dust 2 | 7.46 | 1.394 | 5.36 | <0.001 |
| Soot 0.5 | 0.82 | 0.965 | 0.85 | 0.396 |
| Soot 2 | 1.20 | 0.942 | 1.27 | 0.202 |
| MP fibers 0.5 | 0.64 | 0.980 | 0.66 | 0.511 |
| MP fibers 2 | -1.21 | 1.332 | -0.91 | 0.365 |
| MP fragments 0.5 | -0.47 | 1.128 | -0.41 | 0.679 |
| MP fragments 2 | -0.06 | 1.058 | -0.06 | 0.952 |
| Mix 0.5 | 0.70 | 0.978 | 0.71 | 0.475 |
| Mix 2 | 0.91 | 0.962 | 0.95 | 0.344 |
| Mix 8 | 7.34 | 1.359 | 5.40 | <0.001 |

*Notes.* Model formula: Survival ~ treatment + (1 | GlassID); Family: binomial(logit); SE: Standard error.

Table S3. Regression table for the linear model with knownn sampling variance on earthworm weight. The intercept represents the control treatment. Interactions with time represent earthworm growth per week over eight weeks of exposure. Estimates and p-values are for comparisons with the control.

| | Estimate | SE | z | p |
|---|---|---|---|---|
| Intercept(Control) | 0.31 | 0.018 | 17.44 | <0.001 |
| Brake dust 0.5 | -0.04 | 0.026 | -1.37 | 0.169 |
| Brake dust 2 | -0.02 | 0.024 | -1.02 | 0.307 |
| Soot 0.5 | -0.02 | 0.027 | -0.78 | 0.435 |
| Soot 2 | -0.01 | 0.025 | -0.21 | 0.833 |
| MP fibers 0.5 | -0.04 | 0.022 | -1.79 | 0.074 |
| MP fibers 2 | -0.01 | 0.022 | -0.35 | 0.727 |
| MP fragments 0.5 | -0.03 | 0.023 | -1.13 | 0.258 |
| MP fragments 2 | 0.01 | 0.025 | 0.32 | 0.749 |
| Mix 0.5 | -0.01 | 0.027 | -0.43 | 0.666 |
| Mix 2 | -0.02 | 0.024 | -0.85 | 0.395 |
| Mix 8 | -0.03 | 0.028 | -1.07 | 0.285 |
| Week (Growth in control) | 0.05 | 0.004 | 11.04 | <0.001 |
| Brake dust 0.5:Week | -0.04 | 0.006 | -6.67 | <0.001 |
| Brake dust 2:Week | -0.07 | 0.014 | -5.30 | <0.001 |
| Soot 0.5:Week | 0.01 | 0.006 | 1.12 | 0.263 |
| Soot 2:Week | 0.01 | 0.006 | 1.06 | 0.291 |
| MP fibers 0.5:Week | 0.01 | 0.006 | 1.64 | 0.101 |
| MP fibers 2:Week | 0.00 | 0.006 | 0.10 | 0.922 |
| MP fragments 0.5:Week | 0.00 | 0.005 | 0.79 | 0.429 |
| MP fragments 2:Week | 0.00 | 0.006 | 0.39 | 0.695 |
| Mix 0.5:Week | 0.00 | 0.006 | 0.38 | 0.706 |
| Mix 2:Week | -0.06 | 0.005 | -10.31 | <0.001 |
| Mix 8:Week | -0.09 | 0.007 | -12.33 | <0.001 |

*Notes.* Model formula: Weight ~ treatment * week; SE: Standard error.

Table S4. Regression table for the generalized linear model on the number of cocoons produced by earthworms after 8 weeks of exposure to pollutants. Juveniles were given 4 more weeks to hatch before counting. The intercept represents the control treatment.

| | Estimate | SE | z | p |
|---|---|---|---|---|
| Intercept(Control) | 3.69 | 0.092 | 40.07 | <0.001 |
| Brake dust 0.5 | -5.88 | 1.007 | -5.84 | <0.001 |
| Brake dust 2 | -5.88 | 1.007 | -5.84 | <0.001 |
| Soot 0.5 | -0.71 | 0.142 | -4.99 | <0.001 |
| Soot 2 | -0.78 | 0.144 | -5.44 | <0.001 |
| MP fibers 0.5 | -0.85 | 0.145 | -5.86 | <0.001 |
| MP fibers 2 | -0.06 | 0.131 | -0.47 | 0.639 |
| MP fragments 0.5 | -0.72 | 0.142 | -5.03 | <0.001 |
| MP fragments 2 | 0.01 | 0.130 | 0.05 | 0.962 |
| Mix 0.5 | -2.21 | 0.206 | -10.74 | <0.001 |
| Mix 2 | -5.07 | 0.717 | -7.08 | <0.001 |
| Mix 8 | -3.37 | 0.324 | -10.41 | <0.001 |

*Notes.* Model formula: Cocoons ~ treatment; Family: nbinom2(log); SE: Standard error.

Table S5. Regression table for the generalized linear model on the number of juveniles produced by earthworms after 8 weeks of exposure to pollutants. Juveniles were given 4 more weeks to hatch before counting. The intercept represents the control treatment.

| | Estimate | SE | z | p |
|---|---|---|---|---|
| Intercept(Control) | 3.13 | 0.181 | 17.33 | <0.001 |
| Brake dust 0.5 | -5.33 | 1.028 | -5.18 | <0.001 |
| Brake dust 2 | -5.33 | 1.028 | -5.18 | <0.001 |
| Soot 0.5 | -5.33 | 1.028 | -5.18 | <0.001 |
| Soot 2 | -4.52 | 0.748 | -6.04 | <0.001 |
| MP fibers 0.5 | -4.11 | 0.627 | -6.56 | <0.001 |
| MP fibers 2 | 0.22 | 0.253 | 0.85 | 0.395 |
| MP fragments 0.5 | -5.33 | 1.028 | -5.18 | <0.001 |
| MP fragments 2 | 0.05 | 0.255 | 0.21 | 0.835 |
| Mix 0.5 | -5.21 | 1.029 | -5.06 | <0.001 |
| Mix 2 | -5.21 | 1.029 | -5.06 | <0.001 |
| Mix 8 | -5.33 | 1.028 | -5.18 | <0.001 |

*Notes.* Model formula: Juveniles ~ treatment; Family: nbinom2(log); SE: Standard error.

Table S6. Regression table for the generalized linear mixed model on catalase (CAT) activity (U/mg protein) in earthworms after 8 weeks of exposure to pollutants. The intercept represents the control treatment.

| | Estimate | SE | z | p |
|---|---|---|---|---|
| Intercept(Control) | 1.99 | 0.116 | 17.13 | <0.001 |
| Brake dust 0.5 | 0.54 | 0.166 | 3.26 | 0.001 |
| Soot 0.5 | 0.31 | 0.155 | 2.01 | 0.045 |
| Soot 2 | 0.26 | 0.165 | 1.57 | 0.116 |
| MP fibers 0.5 | 0.24 | 0.155 | 1.57 | 0.117 |
| MP fibers 2 | 0.12 | 0.158 | 0.74 | 0.458 |
| MP fragments 0.5 | 0.19 | 0.150 | 1.24 | 0.216 |
| MP fragments 2 | 0.39 | 0.159 | 2.45 | 0.014 |
| Mix 0.5 | 0.48 | 0.160 | 2.98 | 0.003 |
| Mix 2 | 0.35 | 0.159 | 2.21 | 0.027 |

*Notes.* Model formula: CAT ~ treatment + (1|GlassID); Family: gamma(log); SE: Standard error.

Table S7. Regression table for the generalized linear mixed model on glutathione S-transferase (GST) activity (nmoll/min/mg protein) in earthworms after 8 weeks of exposure to pollutants. The intercept represents the control treatment.

| | Estimate | SE | z | p |
|---|---|---|---|---|
| Intercept(Control) | 16.80 | 4.613 | 3.64 | <0.001 |
| Brake dust 0.5 | -2.91 | 6.738 | -0.43 | 0.666 |
| Soot 0.5 | 5.37 | 6.625 | 0.81 | 0.418 |
| Soot 2 | 16.25 | 6.738 | 2.41 | 0.016 |
| MP fibers 0.5 | -0.51 | 6.433 | -0.08 | 0.937 |
| MP fibers 2 | -4.81 | 6.433 | -0.75 | 0.455 |
| MP fragments 0.5 | 11.47 | 6.142 | 1.87 | 0.062 |
| MP fragments 2 | -3.27 | 6.524 | -0.50 | 0.616 |
| Mix 0.5 | 15.06 | 6.524 | 2.31 | 0.021 |
| Mix 2 | -4.97 | 6.625 | -0.75 | 0.453 |

*Notes.* Model formula: CAT ~ treatment + (1|GlassID); Family: gaussian(identity); SE: Standard error.

Table S7. Regression table for the generalized linear mixed model on malondialdehyde (MDA) activity (U/mg protein) in earthworms after 8 weeks of exposure to pollutants. The intercept represents the control treatment.

| | Estimate | SE | z | p |
|---|---|---|---|---|
| Intercept(Control) | 55.33 | 4.527 | 12.22 | <0.001 |
| Brake dust 0.5 | -8.43 | 6.613 | -1.27 | 0.202 |
| Soot 0.5 | -9.65 | 6.502 | -1.48 | 0.138 |
| Soot 2 | -10.90 | 6.613 | -1.65 | 0.099 |
| MP fibers 0.5 | -13.11 | 6.313 | -2.08 | 0.038 |
| MP fibers 2 | -2.21 | 6.403 | -0.35 | 0.730 |
| MP fragments 0.5 | -16.38 | 6.028 | -2.72 | 0.007 |
| MP fragments 2 | -3.99 | 6.403 | -0.62 | 0.533 |
| Mix 0.5 | -6.57 | 6.403 | -1.03 | 0.305 |
| Mix 2 | -9.68 | 6.502 | -1.49 | 0.137 |

*Notes.* Model formula: MDA ~ treatment + (1|GlassID); Family: gaussian(identity); SE: Standard error.

Table S7. Composition of the different soil mixtures used in the chronic toxicity experiment with *Eisenia fetida*. Treatment soils were prepared by weighing the respective amounts of pollutants and soils to reach the targeted % v/v (volume pollutant/ volume soil) concentrations. In order to be able to convert the targeted concentration (% v/v) into weights, the bulk density (g/ml) of the four pollutants was determined by calculating the mean value from the weight measurements of three 0.5 ml samples each. 100 g of soil was determined to correspond to a volume of 110 ml. For each component, the weight (in g) and respective volume (in ml; within brackets) is given. In addition, the concentration in % w/w (weight pollutant/ weight soil) is given for each treatment.

| Treatment | Concentration (% v/v) | Concentration (% w/w) | PS fragments (density 0.47 g/ml) | PS fibers (density 0.08 g/ml) | Break dust (density 1.89 g/ml) | Soot (density 0.14 g/ml) | Soil (density 0.91 g/ml) |
|---|---|---|---|---|---|---|---|
| *Control* | 0 | 0 | - | - | - | - | 100 g (110 ml) |
| *Single stressor* | | | | | | | |
| PS fragments | 0.5 | 0.26 | 0.259 g (0.55 ml) | - | - | - | 99,600 g (109,45 ml) |
| PS fibers | 0.5 | 0.04 | - | 0.044 g (0.55 ml) | - | - | 99,600 g (109,45 ml) |
| Brake dust | 0.5 | 1.04 | - | - | 1.039 g (0.55 ml) | - | 99,600 g (109,45 ml) |
| Soot | 0.5 | 0.08 | - | - | - | 0.077 g (0.55 ml) | 99,600 g (109,45 ml) |
| PS fragments | 2 | 1.05 | 1.034 g (2.2 ml) | - | - | - | 98.098 g (107.8 ml) |
| PS fibers | 2 | 0.18 | - | 0.176 g (2.2 ml) | - | - | 98.098 g (107.8 ml) |
| Brake dust | 2 | 4.24 | - | - | 4.158 g (2.2 ml) | - | 98.098 g (107.8 ml) |
| Soot | 2 | 0.31 | - | - | - | 0.308 g (2.2 ml) | 98.098 g (107.8 ml) |
| *Multiple stressor* | | | | | | | |
| Mix | 0.5 | 0.35 | 0.065 g (0.1375 ml) | 0.010 g (0.1375 ml) | 0.258 g (0.1375 ml) | 0.019 g (0.1375 ml) | 99,600 g (109,45 ml) |
| Mix | 2 | 1.45 | 0.259 g (0.55 ml) | 0.044 g (0.55 ml) | 1.039 g (0.55 ml) | 0.077 g (0.55 ml) | 98.098 g (107.8 ml) |
| Mix | 8 | 6.16 | 1.034 g (2.2 ml) | 0.176 g (2.2 ml) | 4.158 g (2.2 ml) | 0.308 g (2.2 ml) | 92.092 g (101.2 ml) |
